# Potent binding of 2019 novel coronavirus spike protein by a SARS coronavirus-specific human monoclonal antibody

**DOI:** 10.1101/2020.01.28.923011

**Authors:** Xiaolong Tian, Cheng Li, Ailing Huang, Shuai Xia, Sicong Lu, Zhengli Shi, Lu Lu, Shibo Jiang, Zhenlin Yang, Yanling Wu, Tianlei Ying

## Abstract

The newly identified 2019 novel coronavirus (2019-nCoV) has caused more than 800 laboratory-confirmed human infections, including 25 deaths, posing a serious threat to human health. Currently, however, there is no specific antiviral treatment or vaccine. Considering the relatively high identity of receptor binding domain (RBD) in 2019-nCoV and SARS-CoV, it is urgent to assess the cross-reactivity of anti-SARS-CoV antibodies with 2019-nCoV spike protein, which could have important implications for rapid development of vaccines and therapeutic antibodies against 2019-nCoV. Here, we report for the first time that a SARS-CoV-specific human monoclonal antibody, CR3022, could bind potently with 2019-nCoV RBD (KD of 6.3 nM). The epitope of CR3022 does not overlap with the ACE2 binding site within 2019-nCoV RBD. Therefore, CR3022 has the potential to be developed as candidate therapeutics, alone or in combination with other neutralizing antibodies, for the prevention and treatment of 2019-nCoV infections. Interestingly, some of the most potent SARS-CoV-specific neutralizing antibodies (e.g., m396, CR3014) that target the ACE2 binding site of SARS-CoV failed to bind 2019-nCoV spike protein, indicating that the difference in the RBD of SARS-CoV and 2019-nCoV has a critical impact for the cross-reactivity of neutralizing antibodies, and that it is still necessary to develop novel monoclonal antibodies that could bind specifically to 2019-nCoV RBD.

---

Very recently, a novel coronavirus which was temporarily named “2019 novel coronavirus (2019-nCoV)” emerged in Wuhan, China [1]. As of 24 January, 2020, 2019-nCoV has resulted in a total of 830 laboratory-confirmed human infections, including 25 deaths, in 29 provinces and regions in China, and a number of exported cases in other countries (http://www.chinacdc.cn/jkzt/crb/zl/szkb_11803/jszl_11809/202001/t20200124_211411.html). The person-to-person spread has been confirmed by the findings that there were 15 medical care personnel infections and several family transmission cases, indicating the severe infectivity and pathogenicity of this virus. Currently, however, there is no vaccine or effective antiviral treatment against 2019-nCoV infection.

Based on the phylogenetic analysis (GISAID accession no. EPI_ISL_402124) [2], 2019-nCoV belongs to lineage B betacoronavirus and shares high sequence identity with that of bat or human severe acute respiratory syndrome coronavirus-related coronavirus (SARSr-CoV) and bat SARS-like coronavirus (SL-CoV) (Figure 1a). In previous studies, a number of potent monoclonal antibodies against SARS coronavirus (SARS-CoV) have been identified [3-7]. These antibodies target the spike protein (S) of SARS-CoV and SL-CoVs, which is a type I transmembrane glycoprotein and mediates the entrance to human respiratory epithelial cells by interacting with cell surface receptor angiotensin-converting enzyme 2 (ACE2) [8]. More specifically, the 193 amino acid length (N318-V510) receptor binding domain (RBD) within S protein is the critical target for neutralizing antibodies [9]. We predicted the conformation of 2019-nCoV RBD as well as its complex structures with several neutralizing antibodies (Supplementary Methods), and found that the modelling results support the interactions between 2019-nCoV RBD and certain SARS-CoV antibodies (Figure 1b). This could be due to the relatively high identity (73%) of RBD in 2019-nCoV and SARS-CoV (Figure 1c). For instance, residues in RBD of SARS-CoV that make polar interactions with a neutralizing antibody m396 as indicated by the complex crystal structure [10] are invariably conserved in 2019-nCoV RBD (Figure 1d). In the structure of SARS-CoV-RBD-m396, R395 in RBD formed a salt bridge with D95 of m396-VL. Concordantly, the electrostatic interaction was also observed in the model of 2019-nCoV-RBD-m396, forming by R408 (RBD) and D95 (m396-VL). This analysis suggests that some SARS-CoV-specific monoclonal antibodies may be effective in neutralizing 2019-nCoV. In contrast, the interactions between antibody F26G19 [11] or 80R [12] and the RBD in 2019-nCoV decreased significantly due to the lack of salt bridges formed by R426-D56 in SARS-CoV-RBD-F26G19 or D480-R162 in SARS-CoV-RBD-80R, respectively. Furthermore, while most of the 80R-binding residues on the RBD of SARS-CoV are not conserved on RBD of 2019-nCoV (Figure 1c), it is unlikely that the antibody 80R could effectively recognize 2019-nCoV. Therefore, it is urgent to experimentally determine the cross-reactivity of anti-SARS-CoV antibodies with 2019-nCoV spike protein, which could have important implications for rapid development of vaccines and therapeutic antibodies against 2019-nCoV.

**Figure 1.**
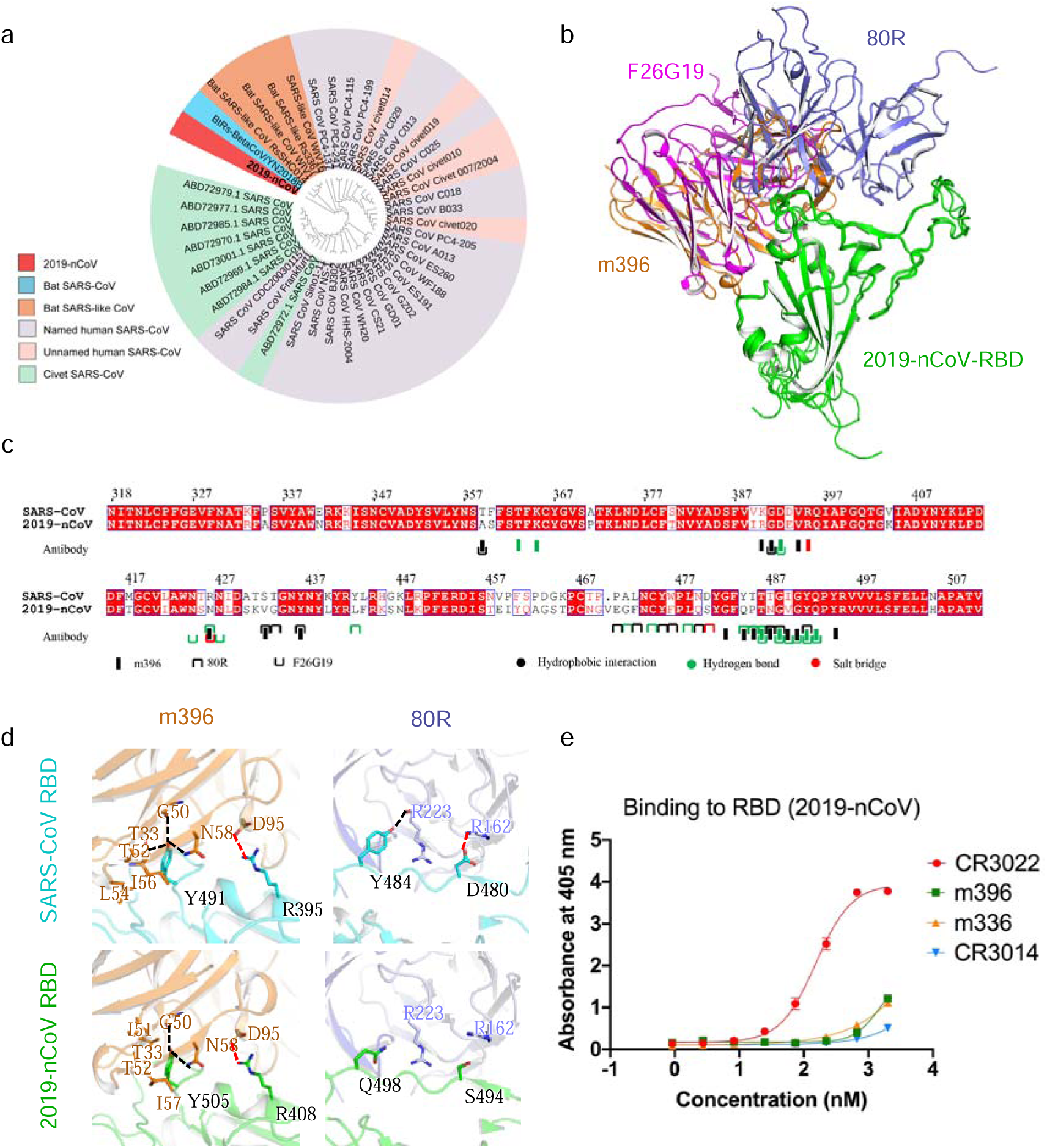
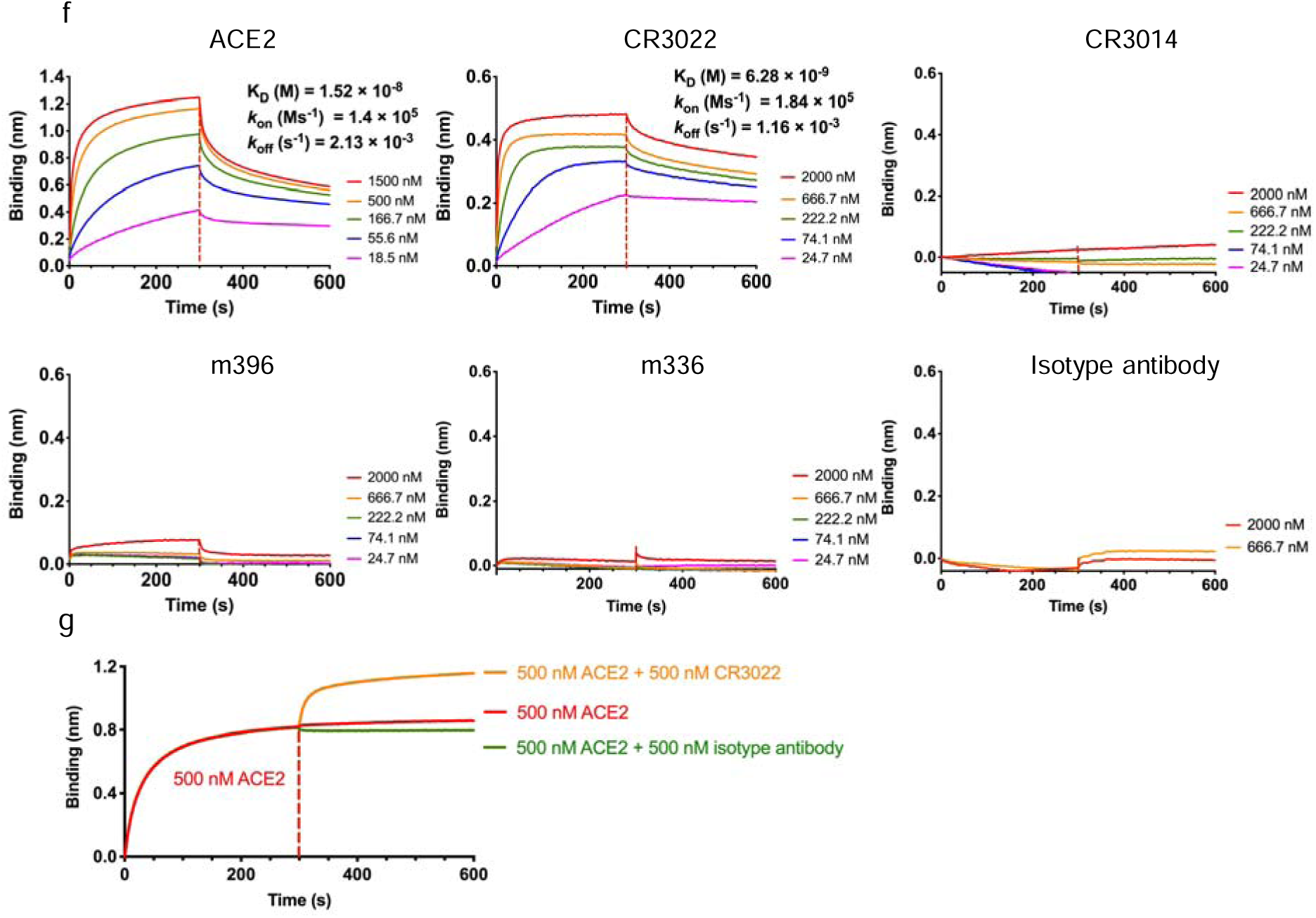
(a) Phylogenetic analysis of 2019-nCoV spike glycoprotein from its protein BLAST sequences. The neighbour-joining tree was constructed using MEGA X, tested by bootstrap method of 2000 replicates, and edited by the online tool of iTOL (v5). (b) The simulated model of 2019-nCoV RBD binding to SARS-CoV-RBD-specific antibodies (m396, 80R, and F26G19). (c) Protein sequence alignment of 2019-nCoV and SARS-CoV RBD, showing the predominant residues that contribute to interactions with SARS-CoV-specific antibodies. (d) The comparison of the complex structures of SARS-CoV-RBD and SARS-CoV-RBD-specific antibodies (shown in the first row) and models of 2019-nCoV-RBD and SARS-CoV-RBD-specific antibodies (shown in the second row). (e) Binding of monoclonal antibodies to 2019-nCoV RBD determined by ELISA. (f) Binding profiles of 2019-nCoV RBD to ACE2 and antibodies, and (g) competition of CR3022 and ACE2 with 2019-nCoV RBD measured by BLI in OctetRED96. Binding kinetics was evaluated using a 1:1 Langmuir binding model by ForteBio Data Analysis 7.0 software.

In this study, we first expressed and purified 2019-nCoV RBD protein (Supplementary Methods). We also predicted the conformations of 2019-nCoV RBD and its complex with the putative receptor, human ACE2 (Supplementary Methods). Comparison of the interaction between complex of ACE2 [13] and SARS-CoV RBD and homology model of ACE2 and 2019-nCoV RBD revealed similar binding modes (data not shown). In both complexes, β5-β6 loop and β6-β7 loop form extensive contact, including at least seven pairs of hydrogen bonds, with the receptor. Notably, R426 on the forth α helix in SARS-CoV RBD builds a salt bridge with E329 and a hydrogen bond with Q325 on ACE2. However, the arginine (R426 in SARS-CoV RBD) to asparagine (N439) mutation in 2019-nCoV RBD abolished the strong polar interactions, which may induce a decrease in the binding affinity between RBD and the receptor. Interestingly, a lysine (K417 in 2019-nCoV RBD) replacement of valine (V404 in SARS-CoV RBD) on β6 formed an extra salt bridge with D30 on ACE2, which may recover the binding ability. These data indicate that the RBD in S protein of 2019-nCoV may bind to ACE2 with the similar affinity as SARS-CoV RBD does. Indeed, we measured the binding of 2019-nCoV RBD to human ACE2 by the biolayer interferometry binding (BLI) assay, and found that 2019-nCoV RBD bound potently to ACE2. The calculated affinity (K_D_) of 2019-nCoV RBD with human ACE2 was 15.2 nM (Figure 1f), which is comparable to that of SARS-CoV spike protein with human ACE2 (15 nM) [14]. These results indicate that ACE2 could be the potential receptor for the new coronavirus, and that the expressed 2019-nCoV RBD protein is functional [2].

Next, we expressed and purified several representative SARS-CoV-specific antibodies (Supplementary Methods) which have been reported to target RBD and possess potent neutralizing activities, including m396 [3], CR3014 [4], CR3022 [5], as well as a MERS-CoV-specific human monoclonal antibody m336 developed by our laboratory [15], and measured their binding ability to 2019-nCoV RBD by ELISA (Supplementary Methods, Figure 1e). Surprisingly, we found that most of these antibodies did not show evident binding to 2019-nCoV RBD. To confirm this result, we further measured the binding kinetics using BLI (Supplementary Methods). An irrelevant anti-CD40 antibody was used as negative control. Similarly, the antibody m396, which was predicted to bind 2019-nCoV RBD (Figure 1d), only showed slight binding at the highest measured concentration (2 µM). Interestingly, we found that there are several additional asparagine residues in 2019-nCoV RBD as compared with SARS-CoV RBD. For instance, the N439 residue is located in the receptor binding site within 2019-nCoV RBD. It could be hypothesized that Asn-linked glycosylation may occur, rendering the computationally simulated models far from the native structures. Further studies are needed to solve the high-resolution structure of 2019-nCoV RBD and understand why it could not be recognized by these antibodies.

Notably, one SARS-CoV-specific antibody, CR3022, was found to bind potently with 2019-nCoV RBD as determined by ELISA and BLI (Figure 1e, 1f). It followed a fast-on (*k*_on_ of 1.84×10^5^ Ms^-1^) and slow-off (*k*_off_ of 1.16×10^−3^ s^-1^) binding kinetics, resulting in a K_D_ of 6.3 nM (Figure 1f). This antibody was isolated from blood of a convalescent SARS patient, and did not compete with the antibody CR3014 for binding to recombinant S protein [5]. To further elucidate the binding epitopes of CR3022, we measured the competition of CR3022 and human ACE2 for the binding to 2019-nCoV RBD (Supplementary Methods). The streptavidin biosensors labelled with biotinylated 2019-nCoV RBD were saturated with human ACE2 in solution, followed by the addition of the test antibodies in the presence of ACE2. As shown in Figure 1g, the antibody CR3022 did not show any competition with ACE2 for the binding to 2019-nCoV RBD. These results suggest that CR3022, distinct from the other two SARS-CoV antibodies, recognizes an epitope that does not overlap with the ACE2 binding site of 2019-nCoV RBD.

The RBD of 2019-nCoV differ largely from the SARS-CoV at the C-terminus residues (Figure 1c). Our results implied that such difference did not result in drastic changes in the capability to engage the ACE2 receptor, but had a critical impact for the cross-reactivity of neutralizing antibodies. Some of the most potent SARS-CoV-specific neutralizing antibodies (e.g., m396, CR3014) that target the receptor binding site of SARS-CoV failed to bind 2019-nCoV spike protein, indicating that it is necessary to develop novel monoclonal antibodies that could bind specifically to 2019-nCoV RBD. Interestingly, it was reported that the antibody CR3022 could neutralize the antibody CR3014 escape viruses, and that no escape variants could be generated with CR3022 [5]. Furthermore, the mixture of CR3022 and CR3014 neutralized SARS-CoV in a synergistic fashion by recognizing different epitopes on RBD [5]. Therefore, CR3022 has the potential to be developed as candidate therapeutics, alone or in combination with other neutralizing antibodies, for the prevention and treatment of 2019-nCoV infections. We expect more cross-reactive antibodies against 2019-nCoV and SARS-CoV or other coronaviruses to be identified soon, facilitating the development of effective antiviral therapeutics and vaccines.

## Acknowledgments

This work was supported by grants from the National Natural Science Foundation of China (81822027, 81630090), National Key R&D Program of China (2019YFA0904400), National Megaprojects of China for Major Infectious Diseases (2018ZX10301403), and the grant from Chinese Academy of Medical Sciences (2019PT350002).

## Declaration of interest statement

No potential conflict of interest was reported by the authors.

