## Supplementary Methods for "Potent binding of 2019 novel coronavirus spike protein by a SARS coronavirus-specific human monoclonal antibody"

***Prediction of the conformation of 2019-nCoV RBD and its complexes with ACE2 or SARS-CoV-RBD specific antibodies***

The homology models of 2019-nCoV RBD domain and its complexes with other components were built on the SWISS-MODEL online server, followed by optimization using Protein Preparation Wizard in Maestro (Schrödinger LLC, New York, NY, USA). The models of 2019-nCoV RBD and its complex with ACE2 were prepared by taking the complex structure of SARS-CoV RBD and ACE2 (PDB entry 2AJF) as template, while the binding modes of 2019-nCoV RBD and the scFv region of SARS-CoV RBD recognizing antibodies m396 (PDB entry 2DD8), 80R (PDB entry 2GHW) or F26G19 (PDB entry 3BGF) were predicted by taking the antibody-antigen complex structures as templates, respectively.

***Protein expression and purification***

The gene encoding RBD (residue 319-541) of 2019-nCoV was synthesized by Genewiz (Suzhou, China), cloned into vector pSecTag containing an N-terminal murine Igκ chain leader sequence and C-terminal His_6_ and Avi Tag. The vector was then transfected into HEK Expi293 cells and incubated at 37°C for 5 days. Supernatant was obtained by centrifugation at 1000 rpm for 10 min and loaded over Ni-NTA as manual described (GE Healthcare, Wayne, PA, USA), washed with wash buffer [10 mM Na_2_HPO_4_, 10 mM NaH_2_PO_4_ (pH 7.4), 500 mM NaCl and 20 mM imidazole] and eluted with elution buffer [10 mM Na_2_HPO_4_, 10 mM NaH_2_PO_4_ (pH 7.4), 500 mM NaCl and 250 mM imidazole]. The collected pure fractions were concentrated and diluted by 10 times of volume of PBS. The procedure was repeated by three times to finally remove imidazole and to replace the storing buffer to PBS. Protein purity was examined with SDS-PAGE, and protein concentration was measured spectrophotometrically at 280 nm.

The heavy chain and light chain sequences of SARS-CoV-specific antibodies m396, CR3014 and CR3022 were connected by a (G_4_S)_3_ linker and subcloned into pComb3x, a vector containing C-terminal His_6_ and Flag tag, and N-terminal periplasmic secretion signal sequence. The antibodies were expressed in *Escherichia coli* HB2151 cells at 30°C for 15 h accompanied with 100 μM IPTG. The cells were harvested and lysed by Polymyxin B at 30°C for 1 h. Protein in supernatant was purified by Ni-NTA after centrifugation at 14000 rpm for 30 mins. Protein purity was examined with SDS-PAGE, and protein concentration was measured spectrophotometrically at 280 nm. The MERS-CoV-specific monoclonal antibody m336 and anti-CD40 isotype scFv antibody were previously prepared and stored in our laboratory.

***Enzyme-linked Immunosorbent Assay (ELISA)***

Costar half-area high binding assay plates (Corning #3690) were coated with purified 2019-nCoV RBD at 100 ng/well in PBS overnight at 4°C and blocked with 3% milk powder (w/v) in PBS buffer at 37°C. 3-fold serially diluted antibodies were added and incubated for 1.5 h at 37°C. HRP-conjugated mouse anti-Flag (Sigma-Aldrich) was used for detection. Enzymatic activity was measured with the subsequent addition of substrate ABTS, and signal reading was carried out at 405 nm.

***Biolayer Interferometry Binding Assays (BLI)***

The binding affinities of ACE2 or m396, CR3014, and CR3022 with 2019-nCoV RBD were performed by BLI on an Octet-Red 96 (Molecular Devices LLC, San Jose, CA, USA) using streptavidin-coated biosensors. RBD of 2019-nCoV was enzymatically biotinylated on C-terminal Avi Tag. The experiment followed a four-step sequential assay at 37°C, samples and buffer were applied in 96-well plate. Next, 6 μg/mL RBD diluted with PBST was loaded on biosensors, tips were dipped into 0.02% PBST for 300 s to reach baseline, then incubated with 3-fold serial diluted antibodies for association, and PBST for dissociation. For the competition analysis of CR3022 and ACE2 binding to RBD, after loading and baseline, tips were first dipped into 500 nM ACE2 in PBST for 300 s, then moved into 500 nM ACE2 in addition with 500 nM CR3022, 500 nM ACE2 or 500 nM isotype antibody control of ACE2 for another 300 s. Results were analyzed by ForteBio Data Analysis software.
